# α-Synuclein emerges as a potent regulator of VDAC-facilitated calcium transport

**DOI:** 10.1101/2020.11.15.383729

**Authors:** William M. Rosencrans, Vicente M. Aguilella, Tatiana K. Rostovtseva, Sergey M. Bezrukov

**Author notes:** To whom correspondence should be addressed: Tatiana K. Rostovtseva, Section on Molecular Transport, *Eunice Kennedy Shriver* National Institute of Child Health and Human Development, National Institutes of Health, 9000 Rockville Pike, Bldg. 29B, Room 1G09, Bethesda, MD 20892-0924.

## Abstract

When the Parkinson’s disease (PD) related neuronal protein, alpha-synuclein (αSyn), is added to the reconstituted mitochondrial voltage-dependent anion channel (VDAC), it reversibly and partially blocks VDAC conductance by its acidic C-terminal tail. Using single-molecule electrophysiology of reconstituted VDAC we now demonstrate that, at CaCl_2_ concentrations below 150 mM, αSyn reverses the channel’s selectivity from anionic to cationic. Importantly, we find that the decrease in channel conductance upon its blockage by αSyn is hugely overcompensated by a favorable change in the electrostatic environment for calcium, making the blocked state orders-of-magnitude more selective for calcium and thus increasing its net flux. These findings reveal a new regulatory role of αSyn, with clear implications for both normal calcium signaling and PD-associated mitochondrial dysfunction.

## Introduction

The mitochondrial outer membrane (MOM), traditionally regarded as a barrier between the mitochondrial space and cytosol, also functions as the interface transmitting signals between the tightly controlled mitochondrial inner membrane (MIM) housing the electron transport chain and the rest of the cell. As the most abundant channel protein in the MOM, the Voltage Dependent Anion Channel (VDAC) controls the permeability of the MOM to ions, water soluble solutes, and metabolites (1–3). Calcium ions (Ca^2+^) are no exception: with the absence of Ca^2+^ specific transporter in the MOM, Ca^2+^ must diffuse through VDAC to cross the MOM (4), though this certainly does not exclude Tom40, the channel protein of the translocase of the outer membrane (TOM complex), as another passive Ca^2+^ pathway across MOM (5). Ca^2+^ is well known as a secondary messenger in the mitochondria. Given VDAC’s unique role as the main conduit for the flux of small molecules and metabolites across MOM, there has been considerable interest in the biophysics of Ca^2+^ transport through VDAC and its regulation. For organellular ion channels like VDAC, the quantitative determination of channel properties is best of all achieved via reconstitution of these channel proteins into planar lipid bilayers (6). Gincel et al first directly demonstrated that VDAC reconstituted into planar lipid bilayer in high concentration pure CaCl_2_ solutions is permeable to Ca^2+^ and gates properly under applied voltage (7). Later, the same group (8, 9) suggested a possible Ca^2+^ binding site at the residue E73, now known from the VDAC1 structure to be imbedded within the lipid membrane (10, 11). Notably, E73 residue is not very conserved among VDAC isoforms and different species: mammalian VDAC isoforms 1 and 2 have it, but mammalian VDAC3 along with VDACs from yeast (12) and fungi (13) have a non-charged glutamine Q73 (see discussion in (14)). Later, Tan and Colombini confirmed the characteristic gating of reconstituted VDAC in CaCl_2_ solutions concluding that channel functions normally with or without Ca^2+^ (15).

Gating, the VDAC’s eponymous phenomenon, is a spontaneous process by which the channel’s unique high conductance “open” state converts into a variety of lower conducting or the so called “closed” states under relatively high applied voltages of > 30 mV (6, 16). Physiological relevance of VDAC gating phenomena is ensured by the fact that its closed states – which are still quite conducting for small ions – are virtually not permeable for ATP (17). Tan and Colombini showed that VDAC closed states, that are typically more cation selective than the open state, have higher permeability to Ca^2+^ (15). By extrapolating their results to physiologically low concentration of Ca^2+^ of 1 μM, they estimated that Ca ^2+^ flux through the closed states could be 4 to 10 times higher than through the anion-selective open state. This study opened the possibility that rather than being an inert pathway for Ca^2+^, VDAC alone or in a complex with other proteins could actively modulate Ca^2+^ signals to or from the mitochondria. Following the work of Tan and Colombini, and taking into account that moving VDAC to its closed states requires application of significant transmembrane voltages, we were interested in learning whether VDAC interactions with protein partners could affect Ca^2+^ flux through the channel.

VDAC has been shown to interact both with other MOM proteins including the cholesterol transporter, TSPO or Bcl-2-family proteins, and various cytosolic proteins such as glycolytic enzymes, hexokinase, dimeric tubulin, and neuronal proteins (18–22). These protein-protein interactions have the potential to modify Ca^2+^ flux through VDAC and thus affect mitochondrial Ca^2+^ homeostasis.

The Parkinson’s Disease (PD) associated protein, α-Synuclein (αSyn) is one of the known potent VDAC regulators (22). It is an intrinsically disordered protein highly expressed in the central nervous system and directly involved in mitochondrial dysfunction in neurodegeneration (23). αSyn is a relatively short 140-residue polypeptide consisting of two distinct parts: a weakly positively charged 95-residue N-terminal lipid-binding domain and highly negatively charged 45-residue C-terminal domain (24). In experiments with VDAC reconstituted into planar membranes it was shown that αSyn is able to block VDAC conductance with nanomolar efficiency (22). The blockage is reversible and partial, with the blocked state still retaining about 40% of the open state conductance. The molecular mechanism is that the negative voltage at the side of αSyn addition drives the negatively charged C-terminal tail of αSyn into the net-positive VDAC pore inducing fast (on the order of milliseconds) characteristic blockages of channel conductance, producing fluctuations of ion current through the channel between its open and well-defined blocked states (25). There are three immediate physiological consequences of this αSyn-VDAC interaction: i) under certain conditions such as a relatively high applied voltage, αSyn can translocate through the VDAC pore and target respiratory complexes in MIM causing impairment of mitochondrial function (26); ii) when the negatively charged C-terminal domain is transiently captured inside the VDAC pore, the blocked state becomes cation selective (27, 28), which should prevent translocation of the negatively charged metabolites (ATP and ADP) due to an electrostatic barrier and steric hindrance; iii) the more cation-selective αSyn-blocked state may suggest that Ca^2+^ permeate more through this state than through the open state of VDAC, which is anionic. The later implication is supported by the data obtained with αSyn overexpression and exogenous addition in HeLa cells where an increase in mitochondrial uptake of Ca^2+^ released by the endoplasmic reticulum (ER) was detected (29). However, direct experiments demonstrating the effect of αSyn on Ca^2+^ flux through the VDAC channel were not available yet.

Another important open question in VDAC physiology is the role of each of the three mammalian VDACs (VDAC1, 2, and 3) in mediating mitochondrial Ca^2+^ signaling. In a key cellular study by De Stefani et al 2012, overexpression or silencing of one of the three mammalian VDAC isoforms was shown to enhance or attenuate the uptake of Ca^2+^ after histamine-evoked release from the ER (30). Overexpression or silencing of VDAC2 or VDAC3, had slightly greater effect on mitochondrial Ca^2+^ uptake than those of VDAC1. Therefore, the authors suggested that this effect may arise from the differences in channel transport properties between isoforms; however, this hypothesis was not explored further. The authors identified that VDAC1 was the only isoform involved in a previously reported mitochondria-ER contact site (MERCS) complex involving the IP3 receptor (IP3R) and Glucose-Regulated Protein 75 (GRP75) (31). This complex was shown to selectively transmit Ca^2+^ signals between the ER and mitochondria. Following the identification of the IP3R-GRP75-VDAC1 complex, other protein complexes with VDAC were found to alter Ca^2+^ signaling, including the Ryanodine Receptor 2 (RyR2) in complex with VDAC2, and most recently the lysosome calcium channel, TRPML1, in complex with VDAC1 (32, 33). Peng et al reported that a mutation in the proposed Ca^2+^ binding site (7), E73Q, reduced Ca^2+^ uptake into the mitochondria.

Here we address the questions of VDAC permeability to Ca^2+^ by utilizing single-molecule electrophysiology on recombinant VDACs reconstituted into planar lipid membranes. We measure ion selectivity of the VDAC open and αSyn-blocked states in CaCl_2_ salt gradient. Using the experimental values of reversal potential and conductance of the VDAC open and αSyn-blocked states in different CaCl_2_ gradients, we calculate Ca^2+^ currents through these states, demonstrating that, notwithstanding channel conductance reduction in the blocked state, αSyn interaction with VDAC actually is able to enhance VDAC-facilitated flux of Ca^2+^ by more than an order of magnitude. We compare the Ca^2+^ selectivity of each individual VDAC isoform as well as of VDAC1 E73Q, where the purported Ca^2+^ binding residue is mutated, demonstrating that while there are significant differences in Ca^2+^ permeability between the isoforms, there is no measurable effect of the E73Q mutation in either the open or αSyn-blocked states. We thus conclude that αSyn interaction with VDAC might have important consequences for Ca^2+^ flux through the pore, while the E73 residue does not seem to play any role in mitochondrial Ca^2+^ uptake. These results have implications for both normal mitochondrial function and PD’s associated dysfunction. It is also worth mentioning that in pursuit of these findings we established that the phenomenon of “charge inversion” is able to take place not only in a single protein channel (34, 35), but at the level of a single polypeptide chain.

## Materials and Methods

### Protein Purification

Recombinant mouse VDAC1 was a kind gift of Dr. Adam Kuszak (NIDDK, NIH). Protocols for expression, refolding, and purification of VDAC1 were based on the work of Dr. Jeff Abramson’s lab (10) as described previously (36). Recombinant mouse VDAC1 E73Q mutant and human VDAC3 were the kind gifts of Jeff Abramson (UCLA). VDAC3 was expressed, refolded, and purified following protocol in (37) and VDAC1 E73Q mutant was purified and refolded as described in (14). All VDAC samples stock solution was 5 mg/ml in 0.1% LDAO. Recombinant WT α-synuclein (αSyn) was the generous gift of Dr. Jennifer Lee (NHBL, NIH). αSyn was expressed, purified, and characterized as described previously (38) and stored at –80°C.

### VDAC reconstitution and conductance measurements

Planar lipid membranes were formed from diphytanoyl-phosphatidylcholine (DPhPC, Avanti Polar Lipids, Alabaster, AL) in 5 mg/ml pentane by apposition of two monolayers of across a ~70 μm aperture in the 15-μm-thick Teflon partition that separates two ~1.5-ml compartments, as previously described (39). Channel insertion was achieved by adding 0.1-2 μL of VDAC sample diluted 100x in buffer containing 10 mM Tris, 50 mM KCl, 1mM EDTA, 15% (v/v) DMSO, 2.5% (v/v) Triton X-100, pH 7.0 to the aperture on the *cis* compartment side of the membrane. Potential is defined as positive when it is greater at the side of VDAC addition (*cis* side). Current recordings were performed as described previously (39) using an Axopatch 200B amplifier (Axon Instruments, Inc.) in the voltage-clamp mode. Unless otherwise noted, single-channel data were filtered by a low pass 8-pole Butterworth filter (Model 900 Frequency Active Filter, Frequency Devices, Inc.) at 15 kHz and saved with a sampling frequency of 50 kHz and analyzed using pClamp 10.7 software (Axon Instruments, Inc.).

VDAC ion selectivity was measured in different CaCl_2_ gradients and in 300 mM (*cis*) / 60 mM KCl gradient buffered with 5 mM HEPES at pH 7.4, as described previously (40). In all experiments VDAC was always added to the *cis* (grounded) compartment. VDAC ion selectivity was inferred from the potential corresponding to the intersection of the current-voltage plot with the zero-current level, the reversal potential. Open state reversal potential was calculated for single and multichannel reconstitutions by applying either a 30 mHz or 50 mHz triangular wave of amplitude 60 mV and/or by measuring conductance acquired at different applied voltages typically at 5 mV intervals. Measurements of the αSyn-blocked state selectivity were carried out only for cases where a single channel was reconstituted. After measuring open state conductance and selectivity, αSyn was added to the membrane-bathing solution to the final 50 nM concentration to the side of higher salt concentration, or both compartments. Data were acquired at different applied voltages after 10-20 min after αSyn addition. Data for measurements of the open state were filtered with low pass 8-pole Bessel filter at 100Hz for open channel selectivity and the higher 500 Hz for the αSyn-blocked state in order to avoid loss of fast αSyn blockage events. Conductance of the open and αSyn-blocked states were determined by the mean of a gaussian fit carried out in ClampFit 10.7. In all cases the current-voltage data were fit with a linear regression to determine both conductance and reversal potential. The measured reversal potential was corrected by the liquid junction potential calculated from Henderson’s equation (34) to obtain the final reversal potential *Ѱ*_*rev*_. The permeability ratios between Cl^−^ and Ca^2+^, *P*_*Cl*_ /*P*_*Ca*_ and between Cl^−^ and K^+^, *P*_*Cl*_ /*P*_*K*_ were calculated according to the Goldman–Hodgkin–Katz equation (41):

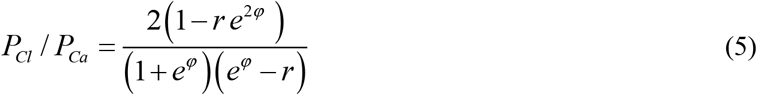

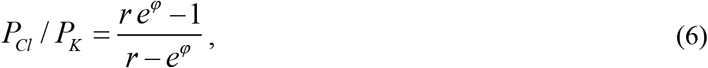

where *r* = *a*_*cis*_ /*a*_*trans*_; *φ* = *Ψ*_*rev*_ (*F*/*RT*), where *R*, *T*, and *F* have their usual meaning of universal gas constant, absolute temperature, and Faraday constant. As the *Ψ*_*rev*_ approaches the Nerst potential, the calculated values for *P*_*Cl*_ /*P*_*Ca*_ approach infinity. These outliers, were excluded for the average value for *P*_*Cl*_ /*P*_*Ca*_ shown in Table 1.

**Table 1.**
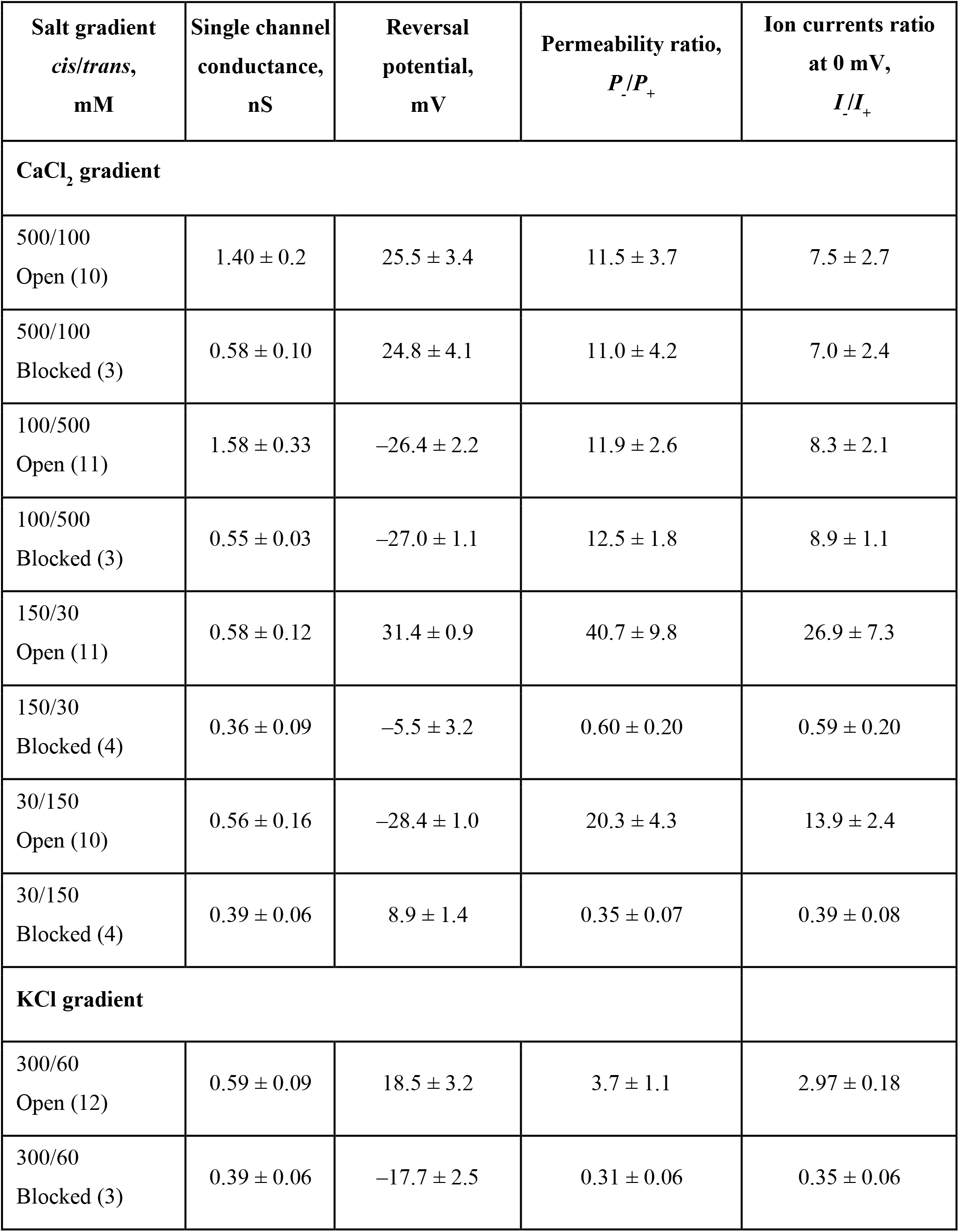
Reversal potential and conductance of VDAC1 open and αSyn-blocked states measured in different CaCl_2_ and KCl gradients. Reversal potential values are corrected for the liquid-junction potential. The ratios of anion to cation permeabilities, *P*_−_/*P*_+_, and currents, *I*_−_/*I*_+_, are calculated according to eq. (5–6) and (4), respectively, at 0 mV. Each data point is a mean ± S.E. Number of independent experiments is shown in brackets. All conditions are as in Figure 3.

| Salt gradient<br>*cis/trans*,<br>mM | Single channel<br>conductance,<br>nS | Reversal<br>potential,<br>mV | Permeability ratio,<br>$P_-/P_+$ | Ion currents ratio<br>at 0 mV,<br>$I_-/I_+$ |
| --- | --- | --- | --- | --- |
| **$\text{CaCl}_2$ gradient** |  |  |  |  |
| 500/100<br>Open (10) | $1.40 \pm 0.2$ | $25.5 \pm 3.4$ | $11.5 \pm 3.7$ | $7.5 \pm 2.7$ |
| 500/100<br>Blocked (3) | $0.58 \pm 0.10$ | $24.8 \pm 4.1$ | $11.0 \pm 4.2$ | $7.0 \pm 2.4$ |
| 100/500<br>Open (11) | $1.58 \pm 0.33$ | $-26.4 \pm 2.2$ | $11.9 \pm 2.6$ | $8.3 \pm 2.1$ |
| 100/500<br>Blocked (3) | $0.55 \pm 0.03$ | $-27.0 \pm 1.1$ | $12.5 \pm 1.8$ | $8.9 \pm 1.1$ |
| 150/30<br>Open (11) | $0.58 \pm 0.12$ | $31.4 \pm 0.9$ | $40.7 \pm 9.8$ | $26.9 \pm 7.3$ |
| 150/30<br>Blocked (4) | $0.36 \pm 0.09$ | $-5.5 \pm 3.2$ | $0.60 \pm 0.20$ | $0.59 \pm 0.20$ |
| 30/150<br>Open (10) | $0.56 \pm 0.16$ | $-28.4 \pm 1.0$ | $20.3 \pm 4.3$ | $13.9 \pm 2.4$ |
| 30/150<br>Blocked (4) | $0.39 \pm 0.06$ | $8.9 \pm 1.4$ | $0.35 \pm 0.07$ | $0.39 \pm 0.08$ |
| **$\text{KCl}$ gradient** |  |  |  |  |
| 300/60<br>Open (12) | $0.59 \pm 0.09$ | $18.5 \pm 3.2$ | $3.7 \pm 1.1$ | $2.97 \pm 0.18$ |
| 300/60<br>Blocked (3) | $0.39 \pm 0.06$ | $-17.7 \pm 2.5$ | $0.31 \pm 0.06$ | $0.35 \pm 0.06$ |

### Statistics

For the statistical analysis of mean values, the difference between two groups of data were analyzed by a one-way ANOVA test using p < 0.05 as the criterion of significance. Differences between many groups were analyzed using the Holm-Sidak multiple comparison tests. All statistical analysis was performed using SigmaPlot. Each experiment was performed a minimum of three times.

## Results

We first verified the previous observations on VDAC channel properties in pure CaCl_2_ solutions (7, 15). Figure 1A shows a representative current record obtained with single VDAC1 channel reconstituted into a planar bilayer formed from diphytanoyl-phosphatidylcholine (DPhPC) that separated 100 mM (*cis*) and 500 mM (*trans*) (100/500) CaCl_2_ solutions (see a schematic of experimental setup in Figure 2A). Under these conditions VDAC1 single channel conductance is 1.6 ± 0.3 nS (mean ± S.D., numbers of experiments are shown in Table 1) and under high applied voltage (−50 mV as in trace in Fig. 1A) the channel moves to a low-conducting or closed state, which is a typical gating behavior of VDAC reconstituted in DPhPC membranes in monovalent salts (42). VDAC characteristic gating in CaCl_2_ gradient could also be monitored under different experimental protocol when a triangular voltage wave of 30 mHz frequency and ± 60 mV amplitude (Fig. 1B, bottom trace) was applied to a single channel inducing a stepwise characteristic channel closure at high voltages (Fig. 1B, upper trace). It can be seen in the Fig. 1A trace that in 100/500 mM CaCl_2_ gradient the reversal potential, *ψ*_*rev*_, (the voltage corresponding to zero current shown in dash-dotted lines in Figs. 1A and 2B) is close to −25 mV (raw data, uncorrected for liquid junction potentials). In the experiment presented in Fig. 1B, where CaCl_2_ gradient was reversed to 500 mM (*cis*) and 100 mM (*trans*) (500/100), the sign of *ψ*_*rev*_ is reversed correspondingly to about +25 mV. Note that VDAC was always inserted from the *cis* side or the grounded side of the planar membrane set up (Fig. 2A). The average data on the reversal potential obtained on at least 10 independent channels in 500/100 and 100/500 CaCl_2_ gradients are shown in Fig. 3A and Table 1. Note that all *ψ*_*rev*_values presented here are corrected for the liquid-junction potentials arising at the KCl salt bridges of Ag/AgCl electrodes, which were calculated using Henderson’s equation (34). Considering that VDAC, as well as other β-barrel channels, such as α-hemolysin (43) or the bacterial porin OmpF (44), is known to predominantly insert into planar membranes in one orientation when added to the same (*cis* in our case) side of the membrane (39), our data show that *ψ*_*rev*_does not depend on channel orientation in 500/100 CaCl_2_ gradient. The corresponding permeability ratios between Cl^−^ and Ca^2+^, *P*_*Cl*_ /*P*_*Ca*_, calculated by eq. (5) are ~11.6 (Fig. 3A and Table 1) for both gradient directions.

**Figure 1.**
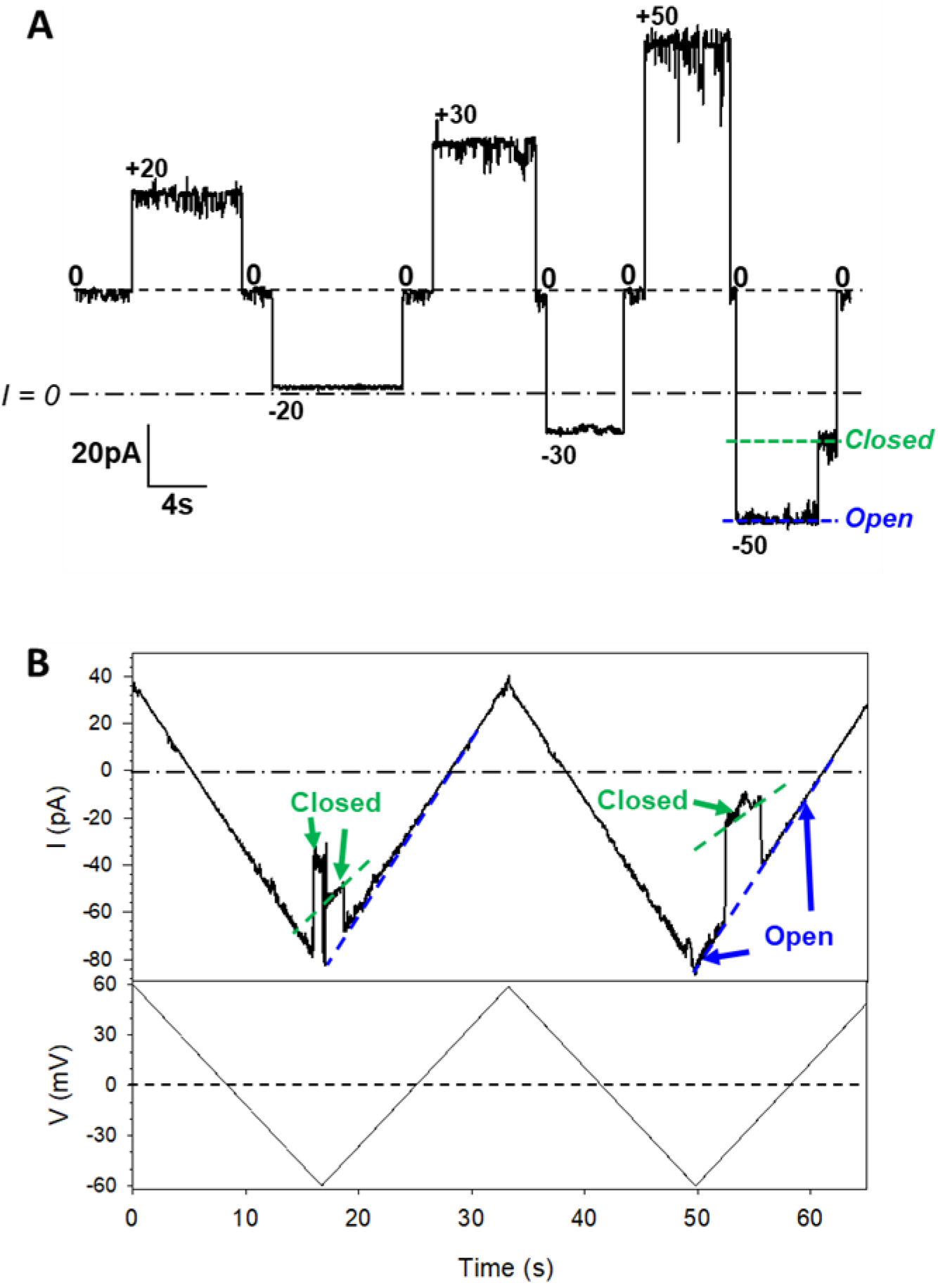
VDAC1 forms typical anion-selective channels in pure CaCl_2_ solutions and characteristically gates under applied voltage. Representative single-channel current traces obtained with reconstituted VDAC1 in 100 mM (*cis*)/500 mM (*trans*) CaCl_2_ gradient at applied voltages as indicated (**A**) and in response to applied triangular voltage wave in reverse 500 mM (*cis*)/100 mM (*trans*) CaCl_2_ gradient (**B**). Applied voltage wave of 30 mHz frequency and ±60 mV amplitude is shown in the bottom panel in (**B**). Here, and elsewhere, the dashed line indicates current at zero applied voltage (V = 0) and the dash-dotted line indicates zero current (I = 0). Note a substantial current at 0 mV in (A). VDAC open and closed states are indicated by dashed blue and green lines, respectively (**A**, **B**). Current records were digitally filtered at 100 Hz using a low-pass Bessel (8-pole) filter. Here, and elsewhere, planar membranes were formed from DPhPC and membrane-bathing solutions were buffered with 5 mM HEPES at pH 7.4.

**Figure 2.**
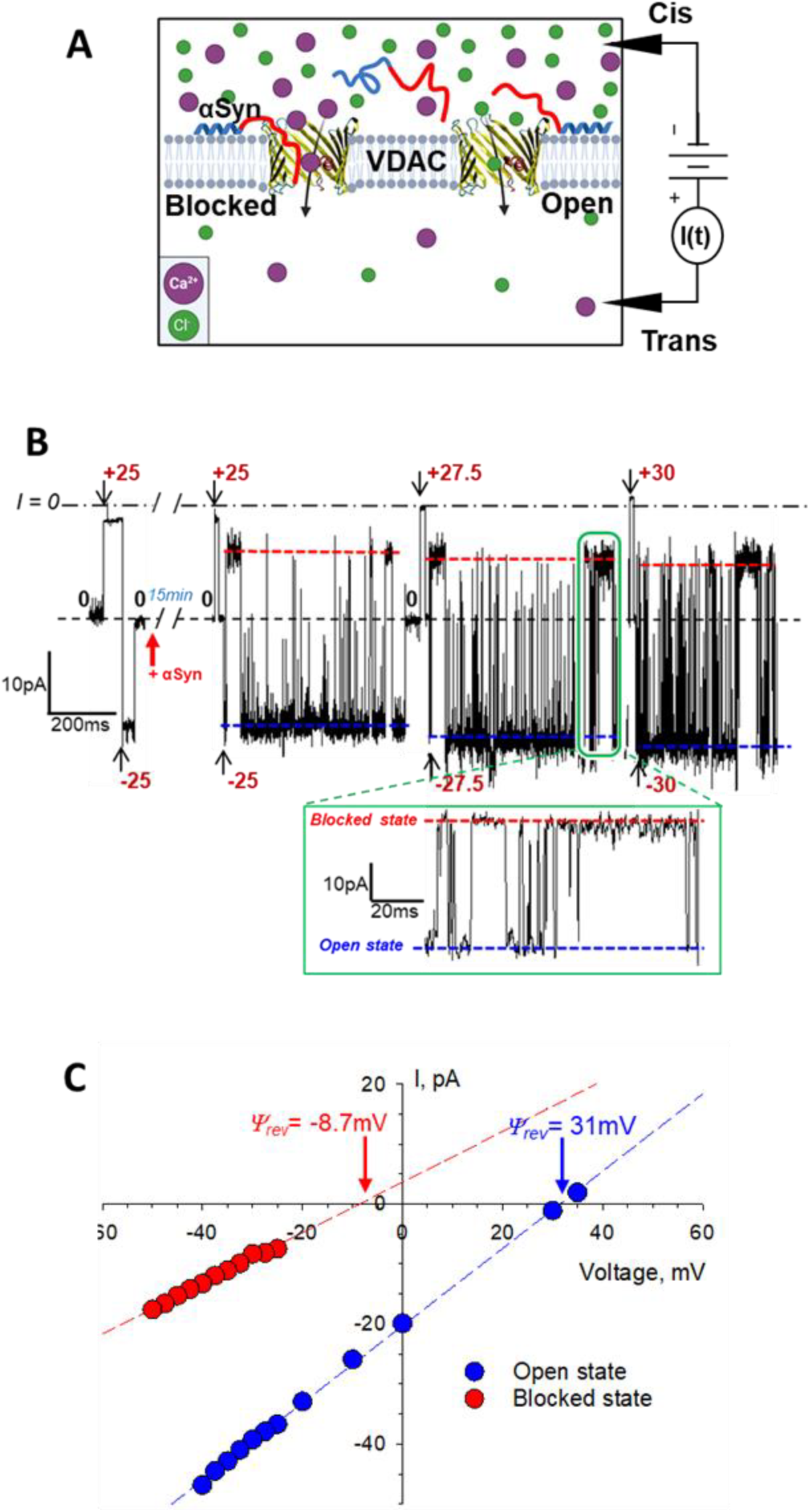
Measurements of ion selectivity of the VDAC open and αSyn-blocked states in CaCl_2_ gradient. **A**– A schematic (not to scale) of experimental setup to measure VDAC ion selectivity with high CaCl_2_ salt in the *cis* and low salt in the *trans* side. αSyn is drawn as a “diblock-copolymer” with the N-terminal membrane binding domain shown in blue and acidic pore-blocking C-terminal domain shown in red. **B**– Representative single-channel trace obtained in 150 mM (*cis*) /30 mM (*trans*) CaCl_2_ gradient before (leftmost trace) and 15 min after addition of 50 nM αSyn to the *cis* compartment at the indicated voltages. Inset shows a fragment of current record at −27.5 mV at a finer time scale. Fast fluctuations of current through the channel between the open and blocked states correspond to individual blockage events induced by αSyn. Blue and red dashed lines indicate the VDAC open and αSyn-blocked states. Current record was digitally filtered at 0.5 kHz using a low-pass Bessel (8-pole) filter. **C**-I/V curves obtained from the traces example of which is shown in (**B**) for the open (blue symbols) and αSyn-blocked (red symbols) VDAC states. Linear regressions (dashed lines) allow calculation of the reversal potential *ψ*_*rev*_(indicated by arrows) for each state. A positive *ψ*_*rev*_corresponds to anion and negative to cation selectivity.

**Figure 3.**
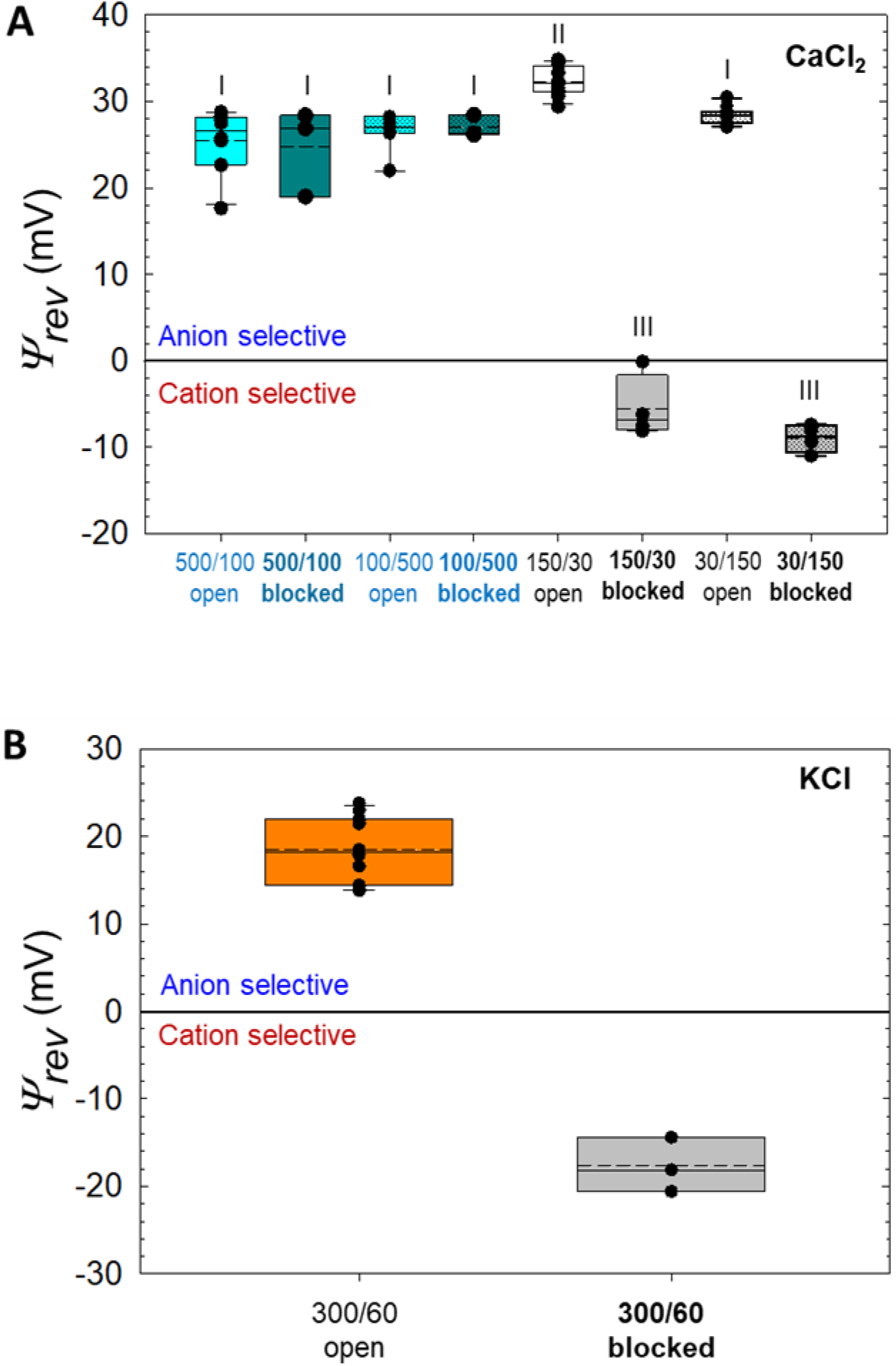
Reversal potentials of the VDAC1 open and αSyn-blocked states measured in CaCl_2_ (**A**) and KCl (**B**) gradients. Salt gradients are shown as in *cis/trans* in mM. Reversal potential (*ψ*_*rev*_) values are corrected for the liquid-junction potential. Note that the sign of *ψ*_*rev*_in experiments in (**A**) with the lower CaCl_2_ concentration in the *cis* side (100/500 mM and 30/150 mM) has been reversed for making comparison of |*ψ*_*rev*_| values easier. Here the ends of the boxes define the 25th and 75th percentiles, with a black line at the median, the mean displayed as dashed black lines, and error bars defining the 10th and 90th percentile. Error bars indicate SD with the numbers of experiments for each condition shown in Table 1. Each different Roman numeral indicates a potential that is significantly different (P < 0.05) from other marked potentials as determined via one-way ANOVA with Holm-Sidak multiple comparison testing. Potentials sharing the same numeral do not differ significantly.

Next, we measured VDAC1 open state selectivity in reduced CaCl_2_ concentration of 150 mM *cis*/ 30 mM *trans* (150/30) and in the reverse gradient of 30 mM *cis*/ 150 mM *trans* CaCl_2_ (30/150). As expected from the previous studies of VDAC selectivity for monovalent ions (45–47), selectivity is more anionic in low CaCl_2_ concentration than in high salt and corresponds to 4-fold higher *P*_*Cl*_ /*P*_*Ca*_ (40.7 ± 9.8) in 150/30 CaCl_2_ gradient than in 500/100 (Fig. 3A and Table 1). Similarly to the previous findings with OmpF (34), in low salt concentrations the selectivity depends on channel orientation. Specifically, in 30/150 CaCl_2_ gradient it is *P*_*Cl*_ /*P*_*Ca*_ = 20.3 ± 4.3 (Fig. 3A and Table 1), only half of that in the opposite orientation. These results are in accord with the earlier data obtained for VDAC in CaCl_2_ solutions (7, 15), confirming that first, VDAC1 gates normally and, second, its open state is anion selective.

Previously we showed that the VDAC αSyn-blocked state is either cationic or non-selective depending on which part of the αSyn molecule is trapped inside the pore at a given time (27, 28). On average, the αSyn-blocked state is essentially non-selective with *P*_*Cl*_ /*P*_*K*_ = 1.0 ± 0.2 measured in 1.0 M *cis* / 0.2 M *trans* KCl gradient (22). Such a reduction in anionic selectivity and in conductance (blocked state conductance is ~40% of the open state) creates strong steric and weak electrostatic barriers for ATP translocation through the VDAC pore suggesting significant reduction or complete abolishment of the transport of large negatively charged metabolites through VDAC due to αSyn-VDAC interaction. Ca^2+^ is a small ion in comparison with ATP or ADP and it might be that even a considerably reduced conductance of the blocked state is still sufficiently large to allow significant Ca^2+^ flux through the blocked pore.

To investigate Ca^2+^ permeability through the αSyn-blocked state, we measured ion selectivity of the blocked state in CaCl_2_ gradients. The schematic of these experiments is shown in Fig. 2A. 50 nM of αSyn was always added to the side with higher CaCl_2_ concentration. A representative experiment with VDAC1 reconstituted in 150/30 mM CaCl_2_ is shown in Fig. 2B. The current records were taken on the same single VDAC1 channel before (*left trace* in B) and 15 min after (*right trace* in B) αSyn addition to the *cis* compartment (Fig. 1A). αSyn induces characteristic time-resolved fast current fluctuations between the unaffected open state and the blocked state, which could be seen best in the inset with a finer time scale in Fig. 2B. Different voltages were applied as shown in Fig. 2B to obtain the corresponding I/V (current-voltage) plots for the open and blocked states (Fig. 2C). The anionic selectivity of the open state quantified as *ψ*_*rev*_= 31 mV is reversed to cationic in the blocked state with *ψ*_*rev*_= −8.7 mV. This corresponds to a change of the average permeability ratio *P*_*Cl*_ /*P*_*Ca*_ from 40.7 ± 9.8 for the open state to 0.58 ± 0.24 for the blocked state (Fig. 3A and Table1), which is the 70-fold change in favor of Ca^2+^permeability. The average *ψ*_*rev*_values, obtained in two CaCl_2_ salt gradients, each in two *cis/trans* configurations, are presented in Fig. 3A and Table 1. *ψ*_*rev*_values for the VDAC open and αSyn-blocked states measured in monovalent KCl salt of the same as in CaCl_2_ 5-fold gradient: 300 mM *cis* /60 mM *trans* KCl, are shown for comparison in Fig. 3B. In KCl gradient, *ψ*_*rev*_is reversed from 18.5 ± 3.2 mV in open to −17.7 ± 2.5 mV in the blocked state with the corresponding ratios *P*_*Cl*_/*P*_*K*+_ of 3.7 ± 1.11 and 0.31 ± 0.06 for the open and blocked states, respectively (Table 1).

Taken together, these results show that in low salt and at small transmembrane voltages the αSyn-blocked state is significantly more permeable for Ca^2+^ than the open state. However, in high CaCl_2_ salt, selectivity of the αSyn-blocked state does not change, remains highly anionic as that of the open state, and does not depend on channel orientation in salt gradient.

Using the same approach, we found that in 150 *cis* /30 *trans* mM CaCl_2_ gradient three mammalian isoforms of VDAC differ significantly in Ca^2+^ permeability. All of them are indeed anionic (Fig. 4A), but with different permeability ratios of *P*_*Cl*_ /*P*_*Ca*_ = 40.7 ± 9.8 for VDAC1, 26.4 ± 11.1 for VDAC2, and 15.0 ± 6.0 for VDAC3. This makes VDAC3 nearly three-fold more permeable to Ca^2+^ than VDAC1, with the Ca^2+^ permeability series of 1 : 1.5 : 2.6 for VDAC1 : VDAC 2 : VDAC 3, respectively. To test the role of the E73 residue as the purported Ca^2+^ binding site, we measured Ca^2+^ selectivity of the VDAC1 E73Q mutant. It turned out that there is no significant difference in Ca^2+^ permeability of the open or αSyn-blocked states between the mutant and WT VDAC1 (Fig. 4B).

**Figure 4.**
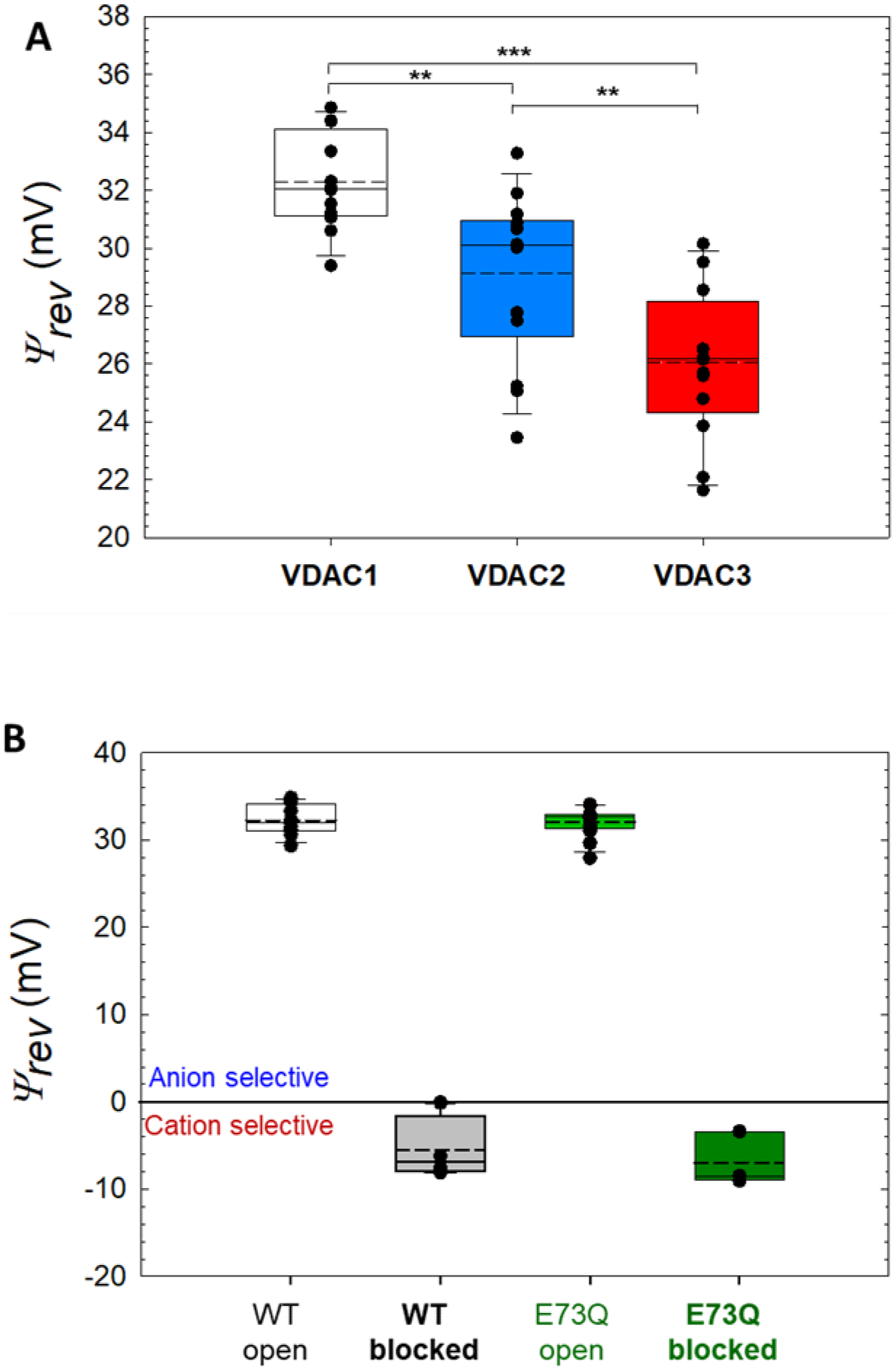
Comparison of ion selectivity of open states for three VDAC isoforms (A) and VDAC1 E73Q mutant (**B**) in 150/30 mM CaCl_2_ gradient. Reversal potentials of the open state of VDAC1, VDAC2, and VDAC3 (**A**) and VDAC1 E73Q mutant (**B**) measured in 150/30 mM CaCl_2_ gradient and corrected for the liquid-junction potentials. Here the ends of the boxes define the 25th and 75th percentiles, with a black line at the median, the mean displayed as dashed black lines, and error bars defining the 10th and 90th percentile. Error bars indicate SD with the numbers of experiments for each condition shown in Table 1. Brackets in (**A**) show significance that was tested using a one-way ANOVA with Holm-Sidak multiple comparison testing (***p < 0.001, **p < 0.01). Differences between data at the open and αSyn-blocked states for WT and E73Q channels in (**B**) are not significant: p > 0.05.

## Discussion

The observed changes in the reversal potential allow us to explore the modulation of VDAC Ca^2+^ permeability by αSyn. Tan and Colombini (15) demonstrated the increase in Ca^2+^ flux induced by VDAC closure. Here, we use a different approach to calculate the total ionic current and its components carried by cations and anions in the open and blocked states from the measured single channel conductance (*G*) and *ψ*_*rev*_. We use the fact that *I/V* curves were linear for the salt gradients of all experiments (see an example in Figure 2C). We also take into account that when the applied voltage equals the Nernst Potential for Ca^2+^ ions, *V*_*Ca*_, their current *I*_*Ca*_ is zero. Analogously, for a bias equal to the Nernst Potential for Cl^−^ ions, *V*_*Cl*_, their current *I*_*Cl*_ is zero. Nernst potential *V*_*i*_ for ionic species *i* with charge number *z*_*i*_ is obtained from ionic activities of the *cis* and *trans* solutions as *V*_*i*_ = – (*RT*/*Fz*_*i*_) ln(*a*_*cis*_/*a*_*trans*_). Total current is *I* = *I*_+_ – *I*_−_; the convention is that *I*_+_ > 0 means cations flowing from *cis* to *trans* while *I*_−_ > 0 means anions flowing from *trans* to *cis*. Then, by considering that *I*(*ψ*_*rev*_) = 0, we get:

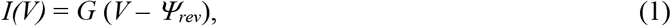

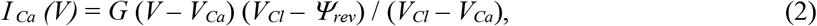

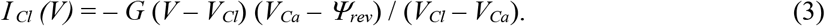

The ratio between anion and cation current at zero voltage becomes independent of the channel conductance and can be expressed as:

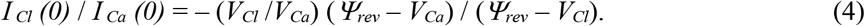

Figure 5 displays individual ion currents through the open (A, C) and αSyn-blocked (B, D) VDAC in 150 mM *cis* /30 mM *trans* (A, B) and 30 mM *cis*/150 mM *trans* (C, D) CaCl_2_ gradients. The actual activity gradient corresponding to 150/30 is 74/19 (48), which yields *V*_*Ca*_ = −17.5 mV and *V*_*Cl*_ = 35.0 mV (both with the opposite sign for the reversed gradient 30/150). The change in the current ratio *I*_*Cl*_ /*I*_*Ca*_ from open to blocked states reflects the change in permeability ratio. In both gradients at zero voltages, *I*_*Cl*_ is an order of magnitude higher than *I*_*Ca*_ in the open state (Figure 5 A, C and Table 1), whereas in the αSyn-blocked state the situation is the opposite: *I*_*Ca*_ is 1.7-2.9 times *I*_*Cl*_ (Figure 5B, D and Table 1). It is seen that the actual *I*_*Cl*_ /*I*_*Ca*_ ratios are indeed voltage-dependent and that gradient inversion gives slightly different absolute values for the ratios. The latter is to be expected (34) due to the structural asymmetry of VDAC.

**Figure 5.**
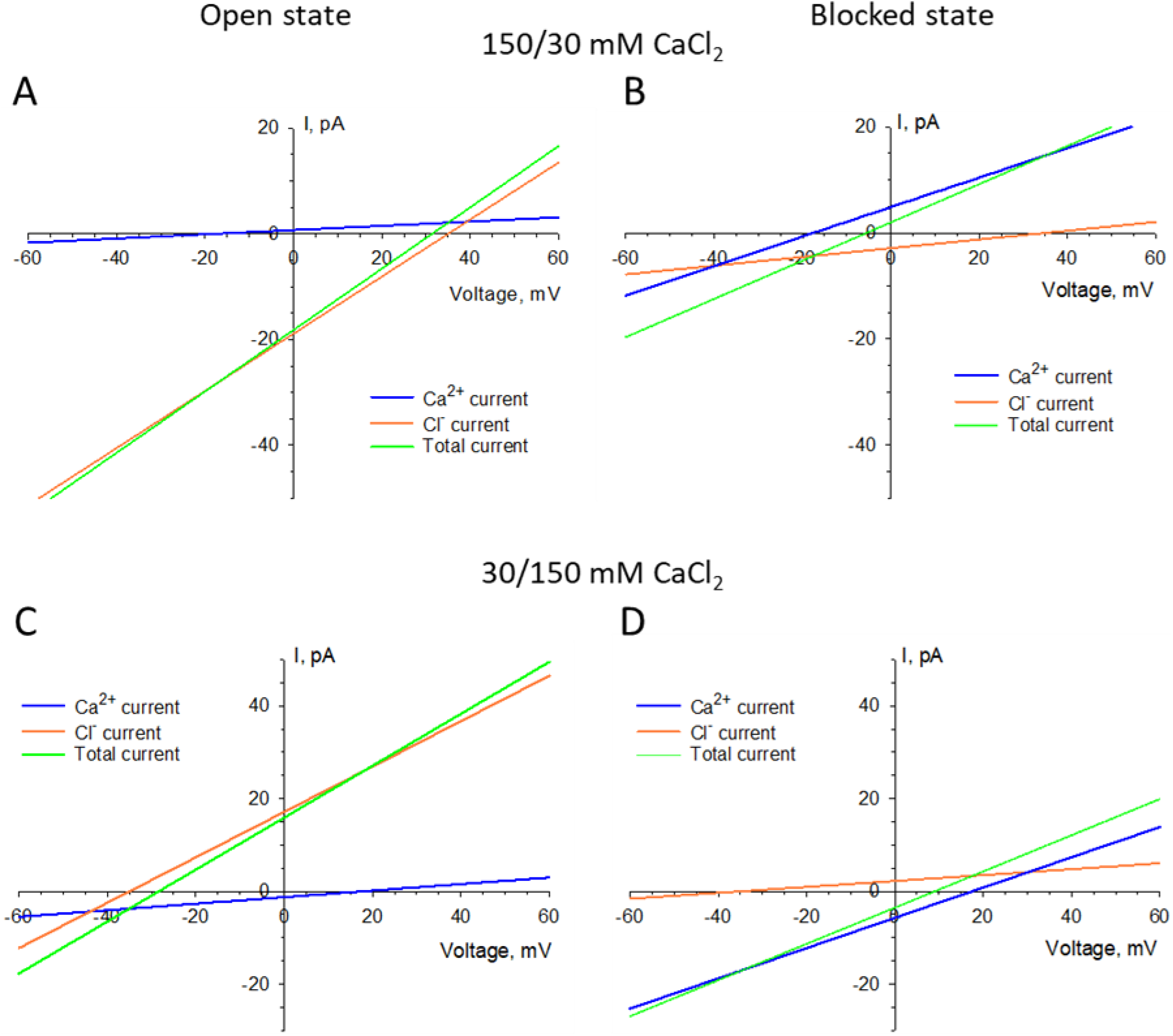
Calculated voltage dependences of individual anionic and cationic components of ion current through the open (**A**, **C**) and αSyn-blocked (**B**, **D**) VDAC in 150 mM *cis* /30 mM *trans* (**A**, **B**) and 30 mM *cis*/150 mM *trans* (**C**, **D**) CaCl_2_ gradients. Currents are calculated based on the average channel conductance and reversal potential (Table 1) according to Eqs. (1–3).

The increase in the Ca^2+^ permeability of the αSyn-blocked state resembles the enhanced Ca^2+^ permeability through VDAC closed states reported by Tan and Colombini (15). Using the 80 mM/ 20 mM CaCl_2_ gradient, they found *P*_*Cl*_ /*P*_*Ca*_ = 24 for the open state. Voltage-induced closure of the channel reduced this value by a factor of 1.5- or 4-fold in favor of Ca^2+^ permeability when channel was moved to the closed states, depending on the “polarity” of the gating. From our experiments in the 150 mM/ 30 mM and in the inversed 30 mM/ 150 mM CaCl_2_ gradient we get the change of more than 50 times in *P*_*Cl*_ /*P*_*Ca*_ in favor of Ca^2+^ permeability upon channel blockage by αSyn (Table 1). Also important, it has to be mentioned here that moving VDAC to its closed state requires much higher voltage biases than those necessary for the efficient interaction with αSyn (22). Although the existence of significant potential differences across the MOM is still under debate (2, 15), the estimates span from 10 mV (49) to as high as 50 mV (50–52), negative at the cytoplasmic side of the MOM. Considering that previous studies suggest that the *cis* side in our bilayer membrane setup corresponds to the cytosolic side of VDAC (53) and that the negative potential applied from the side of αSyn addition is needed to drive the anionic C-terminal tail of αSyn into the pore (22), we can assume that there is a sufficient negative potential across MOM to induce VDAC blockage by αSyn and consequently, increase Ca^2+^ uptake by mitochondria. Indeed, we cannot specify the direction of Ca^2+^ gradient across the MOM *in vivo* because it depends on particular physiological conditions in cell. Nevertheless, calculated *I/V* curves in Fig. 5 allow us to speculate that Ca^2+^ gradient between the cytosol and intermembrane space of mitochondria potentially could either favor mitochondrial Ca^2+^ uptake or its release depending on given physiological conditions.

From the data in Table 1 it follows that there are several ways to describe the main results of this study quantitatively. For example, under the 150/30 mM Ca^+2^ conditions, the blockage can be characterized by the calcium-favoring change (a) in the permeability ratio P-/P+ of 67.8; (b) in the ion current ratio I-/I+ at zero voltage of 45.6; (c) in the current ratio prorated by the conductance reduction of 45.6x(0.36/0.58) = 28.3, and (d) in the permeability ratio prorated for the conductance reduction of 67.8x(0.36/0.58) = 42.08. To add even more complexity, it is necessary to mention that these numbers depend on particular experimental conditions, as clearly demonstrated by Table 1 and Fig. 5. Nevertheless, all the ratios imply that at small concentrations the VDAC-facilitated Ca^+2^ flux is drastically, by more than an order of magnitude, increased by the VDAC interaction with αSyn, notwithstanding the channel conductance reduction in the αSyn-blocked state.

The main goal of the present study was to understand how VDAC interaction with αSyn modifies Ca^2+^ flux through the channel. Indeed, there are two competing factors: (i) an additional steric constraint due to the presence of the αSyn C-terminal tail in the VDAC pore, which reduces its conductance, and (ii) a more favorable electrostatic environment for Ca^2+^ due to the negative charge of the tail. We established that the latter factor dominates, thus increasing the net flux of Ca^2+^. We also showed an interesting effect of the blocked state selectivity reversal by varying Ca^2+^ concentration. Comparison of the data in Fig. 3A and Table 1 shows that the channel is cation selective at the smaller Ca^2+^ concentrations and anion selective at the higher. We relate this to the so-called “charge inversion”, the phenomenon attracting scientists’ vivid attention in many disciplines (54–57). Though it was previously found for a bacterial β-barrel porin, OmpF (34, 35), now, using reconstituted VDAC as a nanopore sensor we demonstrate this phenomenon for a single polypeptide chain – the C-terminal tail of αSyn, the part of the protein that is responsible for the channel selectivity change (28).

One of the immediate implications of these findings to cell physiology is that αSyn could be a potent regulator of mitochondrial Ca^2+^ fluxes by acting through its dynamic interaction with VDAC. Previously reported increase of Ca^2+^ transport from the ER to mitochondria induced by αSyn in permeabilized cells (29) provides support to this conclusion. Notably, the αSyn effect described by Calì et al, required the presence of the C-terminal domain of αSyn. In accord with this observation, our previous data showed that the C-terminal tail of αSyn is a prerequisite for its interaction with the VDAC pore (22). As postulated by Calì et al, WT αSyn may play a physiological role in altering mitochondria Ca^2+^ uptake. This is supported by later evidence suggesting that at physiological levels, αSyn can enhance mitochondrial respiration (58). In contrast, pathogenic expression of αSyn resulting in excess mitochondrial Ca^2+^ uptake may result in mitochondrial dysfunction and cell death. However, one must be extremely careful in direct extrapolations of *in vitro* results to the situation in cell. Genetic evidence is needed to decisively conclude about the role of VDAC in αSyn-induced modulation of Ca^2+^ crosstalk between mitochondria and the ER.

Our experiments with VDAC isoforms suggest that each isoform may differentially mediate Ca^2+^ signaling. VDAC3 was found to be the most permeable to Ca^2+^, followed by VDAC2 and VDAC1. These data are in agreement with De Stefani and coauthor’s hypothesis that slight differences in mitochondrial Ca^2+^ uptake upon overexpression of an individual VDAC isoform are due to differences in isoform Ca^2+^ permeability (30). In their study they found VDAC3 and VDAC2 to enhance Ca^2+^ uptake in comparison to VDAC1. It is tempting to speculate further suggesting that enhanced permeability of VDAC3 to Ca^2+^ may be related to its high expression level in the testes and sperm, where VDAC3 KO in mice leads to male infertility and sperm defects (59). Mitochondrial dynamics and energetics are being increasingly recognized to play an important role in spermatogenesis (60). Likewise, Ca^2+^ signaling plays an important role in various sperm functions including capacitation and the acrosome reaction (61).

The results of the present study are in odds with the statement that the E73 residue of VDAC is a Ca^2+^ binding site (8, 9). VDAC1 structure shows that E73 faces the hydrophobic lipid medium (10, 11) making its accessibility for Ca^2+^ extremely low if not impossible. We recently showed that this residue is not implicated in VDAC1 gating (14). E73 has been found as cholesterol and allopregnanolone binding site (62, 63), and binding of a hydrophobic molecule can influence VDAC gating properties (64). This may give some ground for explanation of why the E73Q mutation in VDAC1 caused a reduction of mitochondrial Ca^2+^ uptake from lysosomes (33).

Our work demonstrates for the first time, that a cytosolic regulator of VDAC modifies this channel properties towards substantially higher Ca^2+^ permeability. Importantly, αSyn is not the only cytosolic protein known to directly interact with the VDAC pore. For example, dimeric tubulin is another known potent VDAC inhibitor (19) which, by blocking the VDAC pore with its C-terminal disordered anionic tail, also reverses VDAC selectivity to cationic (46). This allows us to speculate that tubulin interaction with VDAC could also increase its Ca^2+^permeability, thus opening an intriguing possibility that yet undiscovered VDAC cytosolic protein partners could regulate mitochondrial Ca^2+^ fluxes in an αSyn-like fashion.

## Acknowledgements

W.M.R., T.K.R, and S.M.B were supported by the Intramural Research Program of the National Institutes of Health (NIH), *Eunice Kennedy Shriver* National Institute of Child Health and Human Development (NICHD). V.M.A thanks support from the Government of Spain (PID2019-108434GB-I00 AEI/FEDER, UE), Generalitat Valenciana (AICO/2020/066), and Universitat Jaume I (UJI-B2018-53).

